# Missing interactions: the current state of multispecies connectivity analysis

**DOI:** 10.1101/2021.11.03.466769

**Authors:** Sylvia L.R. Wood, Kyle T. Martins, Véronique Dumais-Lalonde, Olivier Tanguy, Fanny Maure, Annick St. Denis, Bronwyn Rayfield, Amanda E. Martin, Andrew Gonzalez

## Abstract

Designing effective habitat and protected area networks, which sustain species-rich communities is a critical conservation challenge. Recent decades have witnessed the emergence of new computational methods for analyzing and prioritizing the connectivity needs of multiple species. We argue that the goal of multispecies connectivity prioritizations be the *long-term persistence of a set of species* in a landscape and suggest the index of metapopulation capacity as one metric by which to assess and compare the effectiveness of proposed network designs. Here we present a review of the literature based on 77 papers published between 2010 and 2020, in which we assess the current state and recent advances in multispecies connectivity analysis in terrestrial ecosystems. We summarize the four most employed analytical methods, compare their data requirements, and provide an overview of studies comparing results from multiple methods. We explicitly look at approaches for integrating multiple species considerations into reserve design and identify novel approaches being developed to overcome computational and theoretical challenges posed by multispecies connectivity analyses. We conclude that, while advances have been made over the past decade, the field remains nascent in its ability to integrate multiple species interactions into analytical approaches to connectivity. Furthermore, the field is hampered in its ability to provide robust connectivity assessments for lack of a clear definition and goal for multispecies connectivity, as well as a lack of common metrics for their comparison.

## 1. Introduction

Designing effective conservation networks, which sustain species-rich communities across increasingly fragmented landscapes, is a critical challenge for this century as countries commit to the post-2020 Global Biodiversity Framework (Hilty et al. 2020). Central to the success of these networks will be their capacity to meet the connectivity and dispersal requirements of multiple species across remaining habitat areas (Crooks and Sanjayan 2006).

Ecological connectivity measures the extent to which a landscape facilitates or impedes species movement (Crooks and Sanjayan 2006). It is fundamental to species persistence, allowing individuals to seek out food and habitat resources, avoid predation or anthropogenic threats, and promote gene flow (Chetkiewicz et al. 2006; Cushman et al. 2013). A network of connected habitats helps to sustain populations through time (Gonzalez et al. 2011) and to accommodate species undergoing climate or land-use driven range shifts (Pearson and Dawson 2003; Opdam and Wascher 2004; Keeley et al. 2018). Given accelerating rates of habitat loss and climate change, and their negative impacts on animal movement (Tucker et al. 2018), identifying key wildlife corridors that are robust to future environmental conditions is a pressing concern for conservation planners (Crooks and Sanjayan 2006).

Connectivity models are regularly employed to assess habitat networks for individual species (reviewed in Baldwin et al. 2010; Correa Ayram et al. 2016; Arkilanian et al. 2020). However, there is wide consensus amongst scientists and conservation planners for the need to conduct connectivity analyses for multiple species within landscapes. Studies singularly focused on iconic or highly vulnerable species often fail to adequately address the habitat needs of the wider species pool in the landscape (Beier et al. 2009; Cushman and Landguth 2012; DeMatteo et al. 2017; Meurant et al. 2018). Thus, multispecies connectivity (MSC) approaches, which directly or indirectly assess the needs of multiple co-occurring species in a landscape, offer an important avenue to improve spatial conservation planning.

We define an MSC analysis as *“a methodology for identifying a network of habitats and movement pathways that supports the long-term persistence of multiple species in a landscape*”. At a minimum, these analyses must take into consideration connectivity needs of more than one species in a landscape. The ultimate aim of such efforts, however, is to incorporate multiple species interactions into connectivity models and more accurately represent how they mediate habitat use, movement, and the long-term persistence of entire metacommunities (Gonzalez et al. 2011; Chase et al. 2020). This requires moving beyond thinking about connectivity conservation as a “stacking” of habitat networks or metapopulations, towards consideration of multiplex ecological networks in landscapes (Kéfi et al. 2016; Pilosof et al. 2017).

Over the past two decades, various methods have been developed to incorporate the requirements of multiple species into connectivity modelling approaches. Four broad families of approaches have emerged in MSC analyses. The first two integrate multiple species needs at the outset of analysis (hereafter ‘*upstream’* approaches), while the final two integrate them at the end of the analysis (hereafter ‘*downstream*’ approaches):

- *Species agnostic* approaches, such as geodiversity or naturalness methods, which aim to prioritize habitat conservation for multiple species based on the connectivity of biogeoclimatic features and/or the degree to which habitats have been modified by humans (e.g., Koen et al. 2014; Marrec et al. 2020)
- *Generic species* approaches, which combine the traits of multiple species into a single set of values representing the habitat needs and mobility of species groups (e.g. Opdam et al. 2008; Albert et al. 2017);
- *Single surrogate species* approaches, which assess the connectivity requirements of an individual species, selected based on broad habitat needs or sensitivity to disturbance (e.g. umbrella species), to capture the ecological needs of the broader species community (e.g. Brennan et al. 2020); and
- *Multiple focal species* approaches, which separately model connectivity for a set of species representing diverse ecological needs and combine them *post hoc* to identify shared connectivity priorities (e.g., Albert et al. 2017; Meurant et al. 2018; Jennings, Zeller, and Lewison 2020; Williamson et al. 2019).

Currently, there is no general consensus on which of these approaches is most effective for multispecies planning (Marrec et al. 2020). Without a formalized model for implementing multispecies connectivity planning, disparities across methods may have divergent and unintended consequences for conservation design (Reed et al. 2017; Albert et al. 2017; Jennings et al. 2020). MSC assessments can also prove computationally challenging when considering many species in vast landscapes. Identifying faster and less data-intensive approaches with comparable outcomes may be preferred when resources are limited (Santini et al. 2016a) to make MSC assessments more accessible. Given the global push to achieve post-2020 biodiversity goals (IUCN WCPA 2019; Williams et al. 2020), now is a critical time to review progress on multispecies connectivity modelling and operationalize a framework with which countries can achieve their 2050 conservation objectives.

The goal of this study was to evaluate the current state of MSC science in conservation planning. As such, we conducted a literature review to i) assess the frequency of different methods and workflows used to plan for multiple species in connectivity assessments, and ii) evaluate trade-offs across methods in terms of the data and time requirements needed to apply methods and evaluate outputs. We close with a discussion of future directions for this field of research.

## 2. Methods

### 2.1. Literature review criteria

On October 19th, 2020, we used a keyword search in ISI Web of Knowledge scholarly archive to identify scientific articles undertaking multispecies connectivity analyses. We restricted our search to articles published between 2010 and 2020 to focus on the most recent advances in this field. We used the following search terms: ‘Multispecies’ OR ‘Multi-species’ OR ‘Multiple species’ AND ‘Connectivity’ OR ‘Corridor’ OR ‘Surrogate’ OR ‘Geodiversity’ OR ‘Naturalness’ OR ‘Generic species’ OR ‘Focal species’. We also executed a search using the following terms: ‘Connectivity’ AND ‘Focal species’ OR ‘Generic species’ OR ‘Geodiveristy’ OR ‘Naturalness’. Finally, we used ‘Metapopulation capacity’ as a separate search term as it pertains to a new field of research that is related to multispecies connectivity.

From these keywords we identified 503 unique records, which were downloaded along with their abstracts and corresponding publication information. We reviewed abstracts and source journals and restricted the list of articles to i) empirical studies ii) of terrestrial ecosystems, which iii) evaluated connectivity of multiple species, iv) at a landscape-scale or greater. We did however discriminate between single species studies and those that focused on a single umbrella species that was intended to represent the habitat needs and movement requirements of a much larger community of species. After applying these criteria to the abstract of articles, n = 172 papers were retained for in-depth analysis. Articles that met these initial criteria were downloaded and reviewed by the author team. In-depth reading of articles led to the exclusion of an additional n = 95 studies based on the same criteria as mentioned above, leaving n = 77 articles included in the review. Given that some of the reviewed papers explicitly reported on MSC analyses using contrasting methods, we catalogued each of these analyses separately (e.g., Meurant et al. 2018), resulting in 110 case studies (hereafter ‘studies’) of multispecies connectivity modelling for consideration.

### 2.2 Classification and summary of MSC methods

We classified methods reported in the reviewed papers using Arkilanian’s et al. (2020) classification system for connectivity analyses (see Box 1). In this system, connectivity analyses are classified according to a common set of methodological steps in their workflow: 1) species selection, 2) identification of species traits, 3) identification of habitat patches, 4) identification of potential movement pathways between habitat patches, and 5) modelling the degree of connectivity between patches (Arkilanian et al. 2020). Multiple methods exist at each step in the workflow with their own data requirements and capacity to integrate multiple species considerations into connectivity modelling. For methods that did not correspond to a predefined approach, we classified them as ‘Other’ and appended a short description to each record. Some MSC studies carry out connectivity analysis for each focal species separately, and only identify common connectivity priorities *post hoc*. We added a final sixth step to Arkilanian’s classification system to classify different methods used to combine connectivity results at the end of the workflow.

For each study, we collected data on the study location, spatial extent, dominant ecosystem type, study taxa, any software mentioned in connection with the approach used and whether the study contrasted multiple connectivity modelling approaches. We tabulated the number of papers using specific methods at each step in the workflow and identified novel methods that could not be easily categorized in our classification system. We ranked methods in terms of their resource requirements and assessed tradeoffs between computational throughput and precision.

## Box 1.

*Common methodological steps in a multispecies connectivity analysis. Modified from* (Arkilanian et al. 2020)

**Multispecies connectivity analysis workflow**

Multispecies connectivity assessments typically follow a six-step workflow to identify priority areas to conserve connectivity across a landscape. At any point along this workflow methods can be adopted to incorporate consideration of multiple species into the assessment process.

*Step 1. Select species*. Which species are included in a connectivity analysis influences subsequent data and modelling requirements. Four broad approaches exist for species selection: i) species agnostic methods which ignore species-specific data to model landscape characteristics, ii) generic species methods which create a virtual species embodying the characteristics of multiple species, iii) single surrogate species methods which select one species to represent the needs of the wider community, or iv) multiple focal species methods which selected a subset species from the larger pool based on important traits (e.g. phylogeny, taxonomy, functionality, inclusivity).

*Step 2. Identify species traits*. How species are represented in the analysis is based on trait data related to habitat needs, life history and dispersal patterns. These can be derived from direct measures in the field or reported in the literature, the creation of ecoprofiles (sensu Opdam 2008), or by using multivariate approaches that reduce multiple species traits into a singular value (e.g., Laitila and Moilanen 2013). In species agnostic studies this step is skipped.

*Step 3. Define habitat*. What size and types of ecosystems species use to carry out critical portions of their life cycle define which parts of the landscape are considered habitat. Ranking of habitat quality and the classification of what constitutes species’ habitat depends on multiple environmental and ecological factors. Habitat definitions can be informed by GPS or telemetry studies, direct observation (e.g., camera traps, bird counts), distribution or mechanical models, expert opinion, and remote sensing/pattern analysis. The determination of discreet habitat patches is skipped in some methods which instead only rely on a relative ranking of habitat quality.

*Step 4. Define movement capacity*. How far a species can travel and how likely it is to cross less hospitable land covers define a species movement in a landscape. This information is used to determine if a species can travel from patch A to patch B of habitat in a particular landscape. This is commonly achieved by taking information on species habitat preferences and transforming it into a resistance layer by taking the inverse of habitat quality (step 3) and the links between patches established based on species’ dispersal capacity. Studies can use statistical methods, rule-based methods, least-cost paths, circuit theory or through linear programming and optimization to determine and weigh these linkages.

*Step 5. Assess connectivity*. Between which habitat patches and along which routes species are most likely to move define the connectivity of the habitat network. A number of metrics are commonly used to estimate connectivity (i.e. likely movement of individuals). These include benefit maps, conductance/current-density maps, cost maps, graph-theory indices, the metapopulation capacity and/or permeability and area-weighted permeability indices from graph theory. Studies often examine multiple metrics of connectivity to identify key habitat and corridors.

*Step 6. Prioritize multispecies networks*. Which parts of the landscape are most important to conserve species connectivity is based on their ability to maintain movement of species and connect important habitat areas. Prioritization can take place both on a single connectivity map or across multiple connectivity maps for different species/groups. Results from multiple connectivity analyses can be combined by assessing areas of spatial overlap amongst connectivity maps, tabulating the number of overlapping species networks across different parts of the landscape, normalizing and summing connectivity values across species, combining the top ranking percentile of connectivity values for each species or through a process of optimization.

## 3. Results

### 3.1. Characterizing MSC studies

Many of the studies initially retained for review based on keywords in the title and abstract did not analyze MSC directly. After applying our selection criteria, 77 papers were retained detailing 110 studies of multispecies connectivity assessments published between 2010 and 2020 (Fig.1A). In most cases, excluded studies alluded to the importance of multiple species considerations in conservation planning and in their selection of a single focal species but provided no further analysis on the relevance of the selected species to the wider community. A second set of studies that were excluded looked at the connectivity needs of multiple species but did not include methods to identify a common connectivity prioritization across species. A third group of studies aimed to predict species’ use of the landscape based on predictors of habitat quality for multiple species and discussed connectivity amongst habitats but did not directly model the movement pathways through the landscape. Over the considered timeframe the annual number of published studies increased gradually with a spike in publications occurring in 2017 and 2019. As the literature search was conducted in October of 2020, it is possible that a number of additional papers were also published at the end of this year that were not considered.

**Figure 1.**
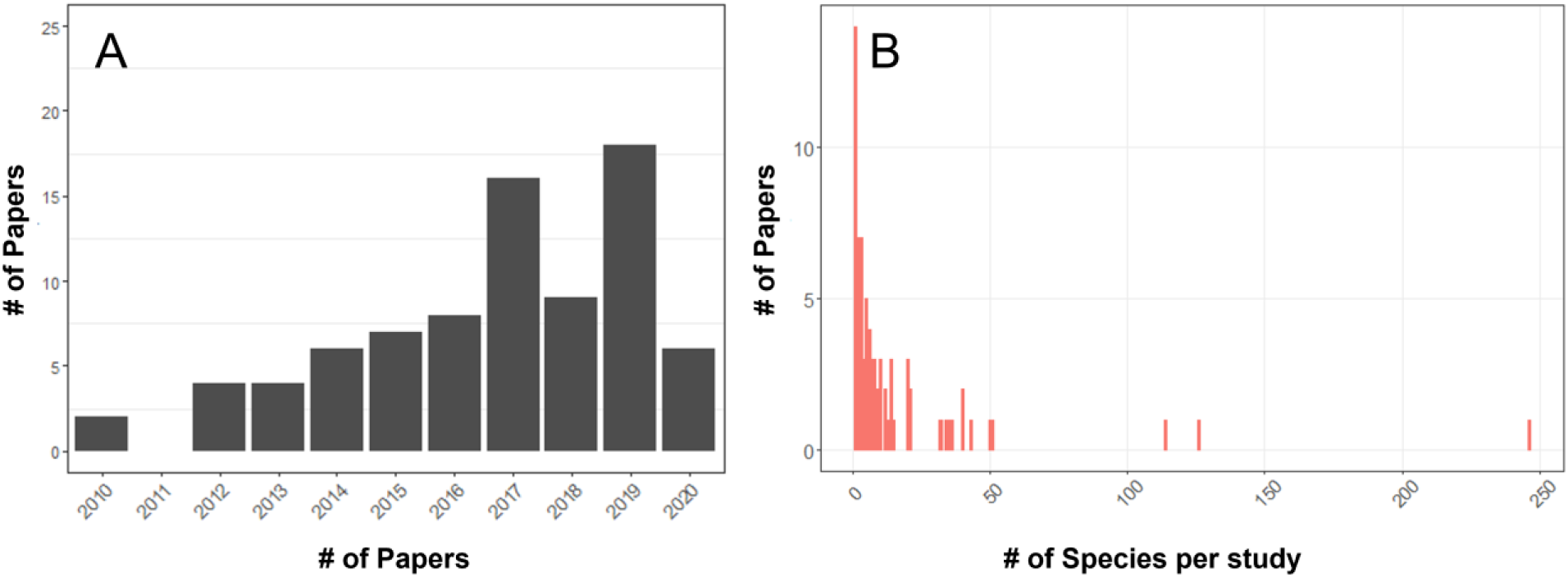
(A) The number of papers meeting the criteria of an inclusion in the literature review between 2010 and 2020 (total n = 77), and (B) the distribution of the number of species assessed per study across reviewed papers (one study excluded with >2000 species)

Most retained studies were located in North America, Western Europe, China, and Australia with a smaller number of studies from Southeast Asia and South America (Fig. 2A). A few supra-national studies were also included in the review which looked at connectivity in regional Austral-Asian flyways (Iwamura et al. 2014), the European Alps (Hanson et al. 2019), as well as global patterns of connectivity amongst protected area networks (Santini et al. 2016b), forests in global biodiversity hotspots (Larrey-Lassalle et al. 2018), and tropical mangrove ecosystems (Huang et al. 2020).

**Figure 2.**
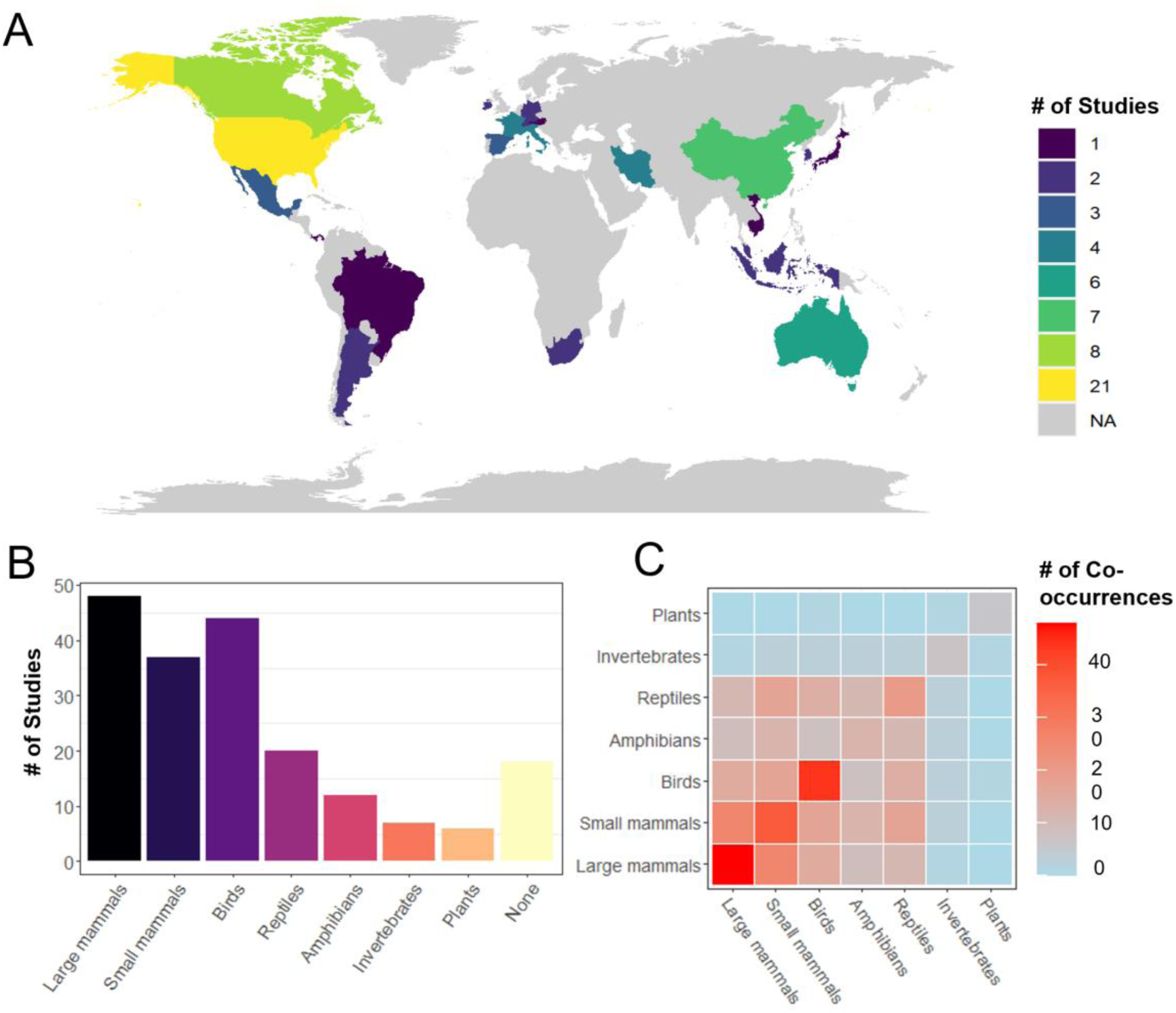
A) Global distribution of reviewed multispecies connectivity studies (total n=110), B) the number of reviewed studies evaluating the connectivity of each considered taxonomic group included or no species (None) as in species agnostic approaches, (C) a heatmap of the number of times species of a taxonomic group were co-assessed with species from the same or other taxonomic groups within studies. Warm colours indicate higher frequencies and cooler colours indicate lower frequencies

In the reviewed papers, the average number of species considered was 16.6 ± 34.5 (min = 0, max = 246, one studied excluded with > 2000 plant species, Fig 1B). Of the 110 studies reviewed, 52% focused on connectivity patterns of multiple species within a single taxonomic group (n = 57, small and large mammals were combined), while 47% (n = 53) looked at species across taxonomic groups (Fig 2C). The most frequently considered taxonomic groups were large mammals (44%) followed by birds (40%), small mammals (34%) and then reptiles (18%) and amphibians (11%), with few studies focused on invertebrates (7%) or plants (5%) (Fig. 1b). Of studies satisfying our criteria, 18% applied a species agnostic approach, 17% took a generic species approach, 9% used a single surrogate species to assess the wider community connectivity, and the remaining 55% took a multiple focal species approach (Table 1). Overall, most reviewed studies assessed connectivity in large landscapes (1,000-10,000 km^2^) up to the scale of subcontinents (>100,000km^2^) (Table 1) and were predominantly focused on temperate forests, agro-ecosystems, or landscapes that encompassed multiple large ecosystem types.

**Table 1.** Frequency of studies employing each of the four broad approaches for species selection per spatial scales of consideration.

| Scale | Extent (km<sup>2</sup>) | Species agnostic | Generic species | Surrogate species | Multiple focal species | Total |
| --- | --- | --- | --- | --- | --- | --- |
| Small landscape | 10-100 | 1 | 1 | 0 | 3 | 5 |
| Medium landscape | 100-1,000 | 2 | 2 | 0 | 8 | 12 |
| Large landscape | 1,000-10,000 | 7 | 6 | 4 | 12 | 29 |
| Ecoregion | 10,000-100,000 | 4 | 6 | 4 | 20 | 34 |
| Sub-continental | >100,000 | 5 | 4 | 2 | 16 | 27 |
| Continental | NA | 1 | 0 | 0 | 2 | 3 |
| **Total** | **-** | **20** | **19** | **10** | **61** | **110** |

### 3.2. Upstream and downstream approaches

Multispecies connectivity approaches invariably require collapsing information on multiple species’ habitat or movement needs at some point in their analytical workflow. This can occur prior to calculating connectivity metrics, what we term *upstream* approaches, resulting in a composite connectivity map from a single connectivity analysis. In contrast, *downstream* approaches build connectivity maps for each individual species and then collapse them to arrive at a composite prioritization map. Some studies can adopt both upstream and downstream approaches in their workflow by collapsing species information into multiple generic species at the outset of analysis and afterwards combining connectivity results from their analysis (e.g., Ecoprofiles, Opdam et al. 2008). One exception to our dichotomization of upstream and downstream approaches is the use of optimization algorithms during the connectivity analysis (step 5) to incorporate multiple species considerations simultaneously (e.g. Wang and Önal 2016).

Upstream and downstream approaches vary in the computational resources required to parameterize connectivity models, and approaches that collapse multiple species information earlier in the workflow are less data-intensive. Upstream methods, such as species agnostic or generic species approaches, require substantially less species-specific data and computational resources than studies that collapse species information in steps 3-5. Upstream approaches are also much less data-intensive than multiple focal species connectivity analyses that requires carrying species-specific data through each step of the workflow. This tradeoff can become increasingly important as the resource requirements increase exponentially as the number of species and landscape extent increase (Santini et al. 2016a).

### 3.3. Common approaches in MSC analyses

We found the majority of reviewed studies applied downstream approaches when analyzing MSC by adopting a multiple focal species approach (Fig. 4). These studies selected a subset of species in their landscape (Fig. 3, step 1) using an array of criteria (e.g., representativeness, functional roles, vulnerability) and proceeded to carry out individual connectivity analysis for each species separately (steps 2-5). They then combined connectivity maps to identify common priority areas for connectivity (step 6), principally through overlap analysis or by summing multiple connectivity layers (Fig. 3, left hand column). A smaller number of studies adopted upstream approaches by adopting a species agnostic or generic species approach for their MSC analysis or combining multiple species data prior to running the connectivity analysis. These approaches are generally less data-intensive and are growing in frequency in more recent years.

**Figure 3.**
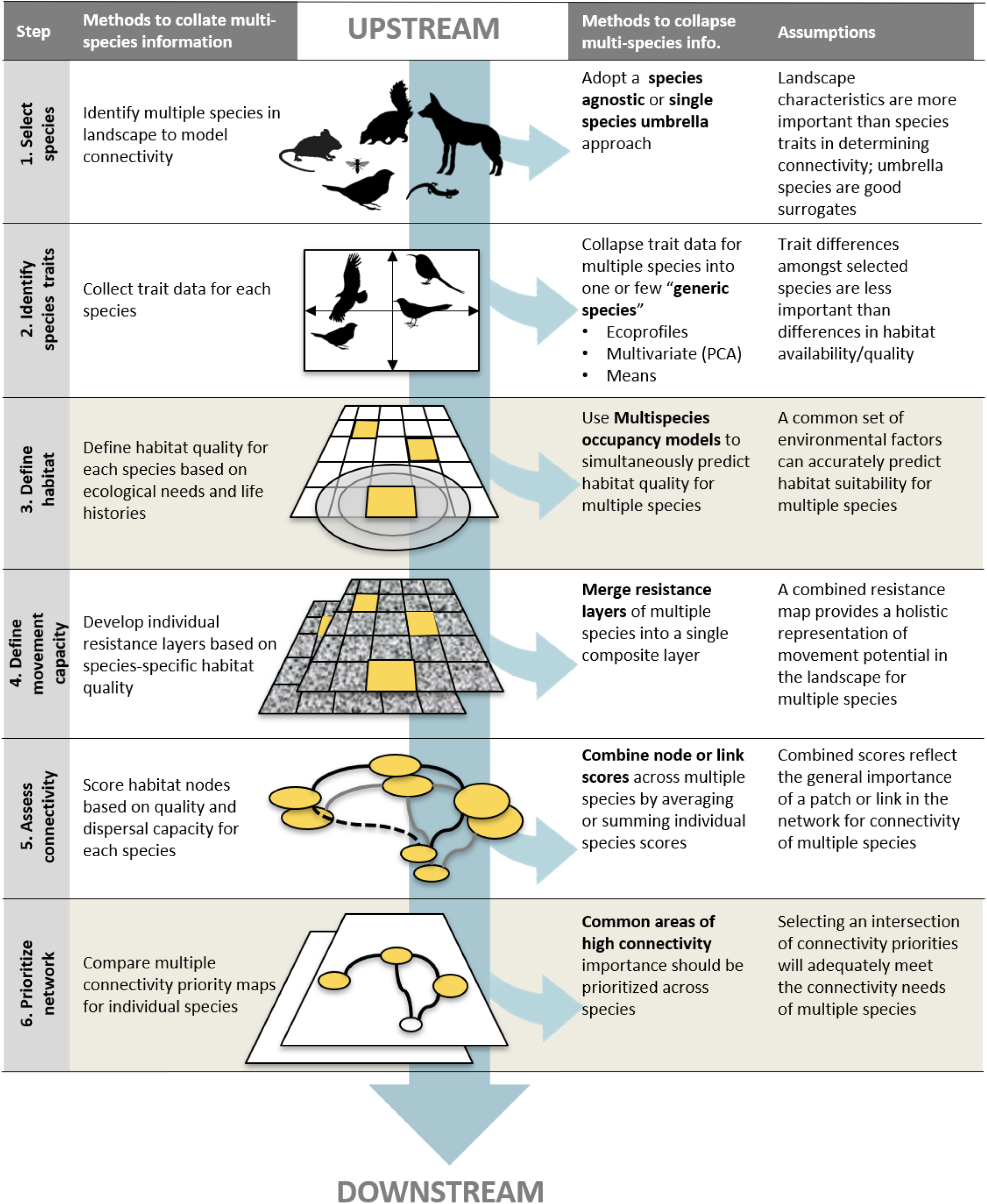
A conceptual diagram of the six-step workflow in multispecies connectivity assessments characterizing upstream vs. downstream approaches to incorporate multiple species information. The column on the left indicates the general analysis followed in a multiple individual species connectivity assessment for each step. The columns on the right identify potential methods for collapsing multiple species information at each step and the associated assumptions. Steps 2 and 6 are shaded to indicate that they are not necessarily included in all multispecies connectivity assessments.

**Figure 4.**
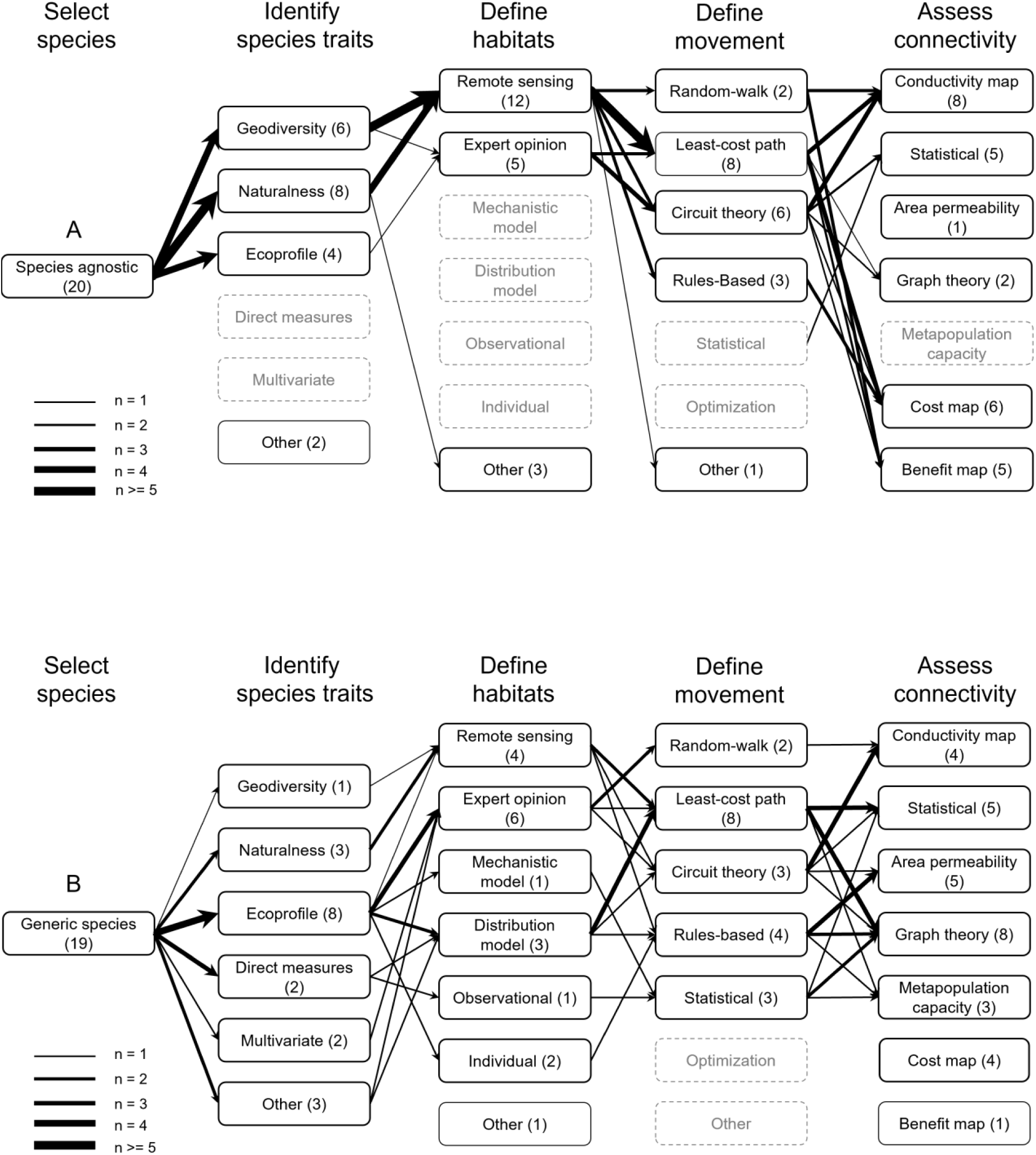
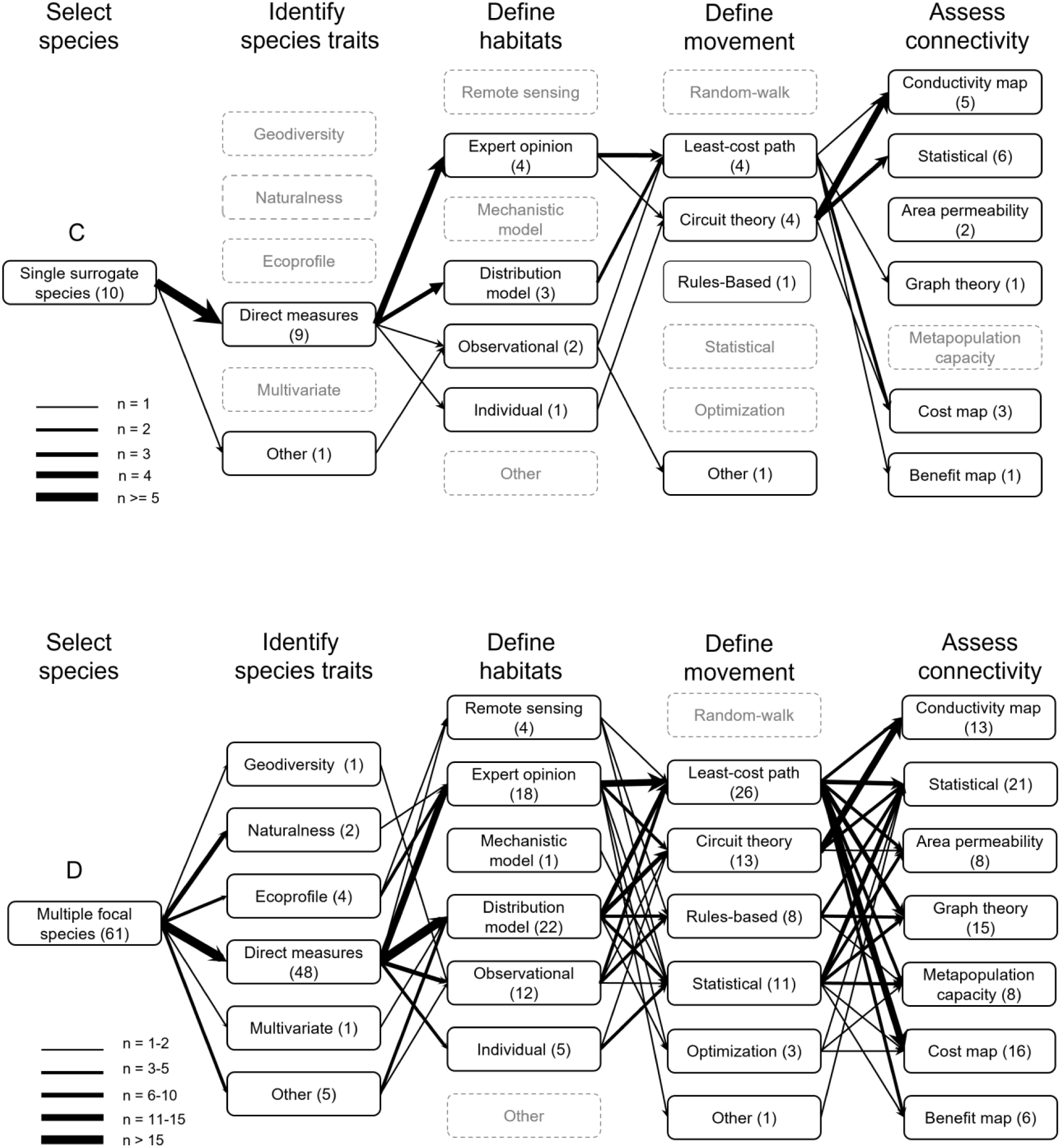
Connectivity workflows for four common multispecies selection approaches: (A) species agnostic, (B) generic species, (C) single surrogate species and (D) multiple focal species. Boxes represent different available methods to employ for each step along the workflow. The number of studies employing each method is indicated in parentheses; the thickness of arrows linking methods across steps shows the frequency of method combinations (see legends). Greyed out boxes indicate methods that were not used by any workflow reviewed.

Across all reviewed studies, the most common methods used to assess potential movement across a landscape (step 4), whether for an individual or generic species, was through least-cost path analysis and/or circuit theory analysis. Most studies also combined multiple methods for their connectivity assessment (step 5), the most common being graph theory metrics with either least-cost or current density maps. A handful of studies also calculated the metapopulation capacity, which is a measure of the capacity of a given landscape configuration to support the persistence of a specific species (Ovaskainen and Hanski 2001). Very few studies used optimization approaches to identify common connectivity priorities amongst multiple species and none explicitly incorporated species interactions into their methods.

### 3.4. Comparing effectiveness of MSC approaches

Opting for simplified upstream approaches may be desirable or necessary in situations of limited data and/or computational capacity. However, given that connectivity maps are sensitive to the way in which connectivity is formalized and implemented in models (Reed et al. 2017; Albert et al. 2017), it is important to understand the potential trade-offs of the different MSC approaches when modelling the connectivity needs of diverse species. In our review we came across eight papers that explicitly compared the results from two or more of the broad classes of MSC approaches, each using a multiple focal species approach as the basis for comparison.

The most frequent comparison was between a single surrogate species and a multiple focal species approach. Across these studies, single surrogate species approaches were found to poorly capture the connectivity and habitat needs of the wider species community (Brodie et al. 2015; DeMatteo et al. 2017; Meurant et al. 2018; Brennan et al. 2020).

Studies comparing generic species and multiple focal species approaches found significant differences in the priority rank-maps of the generic species to the composite maps, but overall connectivity maps of generic species performed better than single surrogate species (Brodie et al. 2015; Meurant et al. 2018). Importantly, however, Brodie et al. (2015) found in Borneo that their generic species approach became increasingly effective as the degree of ecological similarity and/or sensitivity to disturbances increased amongst the represented species in their tropical forest community.

Finally, studies comparing species agnostic approaches, which do not rely on any species-specific data, to multiple focal species approaches have been more varied in performance (Brost and Beier 2010, Koen et al 2014, Jennings et al 2020). The inconsistency of findings both within and amongst species agnostic studies suggest that more work is needed to refine and validate these approaches before they can be used with confidence to capture the needs of diverse communities of species in landscapes.

To date, the few papers comparing methods seem to suggest that single surrogate species models may poorly represent the habitat and connectivity needs of the wider community of species. In contrast, carefully constructed generic or virtual species, where represented species share similar ecological traits, may provide a more promising approach to model multiple species connectivity when data and processing capacity is limited.

Finally, Williamson et al. (2020) compared the impact of different *post hoc* methods to combine connectivity analyses outputs in a multiple focal species approach. Their results underscore the challenge of consolidating multiple aspects of species biology into a single map (Williamson et al. 2020). Each of the methods to combine species connectivity maps had limitations either in their ability to fairly represent the habitat needs and movement capacities of different species (normalized sum), may overlook moderate-to-high value habitats that could support multiple species (top percentile), and/or were sensitive to the selected threshold delimiting habitat from non-habitat (model count). Being aware of the limitations of each with regard to the type of species under consideration will be important to selecting the most appropriate metric to combine multiple connectivity analyses.

### 3.5. Novel methods in MSC

In addition to approaches that incorporate multispecies considerations either at the outset of the analysis (*upstream*) or as a final step in the connectivity analysis (*downstream*), our literature review uncovered several studies that employed novel methods for calculating MSC (see Fig. 3). Starting with a subset of species from the landscape, these studies combined individual species information at different points along the workflow to produce a single connectivity assessment or prioritization.

#### i) Multispecies occupancy models

Multispecies occupancy models can be used to predict species locations and connectivity across landscapes when individual species presence-absence data are scarce (Meyer et al. 2020). This allows for a single model to predict the occupancy of habitat patches for multiple species, some of which may be rare and difficult to detect. In their study, Meyer et al. (2020) used camera-trap data for nine medium to large mammals and a hierarchical multispecies occupancy model to estimate species occupancy in the Mesoamerican Biological Corridor. They estimated species-specific model parameters as random effects of a community-level distribution, which permits more precise parameter estimates for rare species than traditional species-level analyses (Zipkin et al. 2010; Kéry and Royle 2015). From this, the authors developed an occupancy-weighted connectivity metric to evaluate species-specific functional connectivity. While Meyer et al. (2020) stop short of a full multispecies connectivity assessment by not identifying common priority areas of connectivity, their methodology could be used to great effect to improve multispecies habitat identification in data limited contexts.

#### ii) Combining habitat suitability and resistance layers

In their study in central-western Mexico, Correa Ayram et al. (2019) developed common habitat suitability and resistance layers for three multispecies groups to identify composite multispecies corridors. Starting with 40 focal species with contrasting habitat needs, Correa Ayram and colleagues (2019) grouped species based on shared inter-patch dispersal distances and minimum habitat requirements. Within each multispecies group, a common habitat layer was developed by retaining only habitat patches which were common to all species. Additionally, individual species resistance layers were summed and normalized to build a common resistance layer for each multispecies group. These layers were then used as inputs into least-cost path and circuit theory analyses to prioritize commonareas of connectivity importancen. This approach could be considered a variant of Opdam et al.’s (2008) ecoprofile approach, however, by collapsing individual species information after developing and employing species-specific habitat models, Correa Ayram and colleagues carry forward a greater amount of species-specific habitat information along the workflow.

#### iii) Combining node and link metrics

In their study of all non-volant terrestrial mammals in Italy, Santini et al. (2016a) aimed to reduce the computational effort associated with large MSC assessments by combining probabilistic species graphs prior to conducting the network analysis. In their study, the authors tested multiple methods for aggregating node attributes (summing values of the probability of connectivity and of intra-patch connectivity) and link attributes (mean, weighted-means) for all species to increase computational efficiency. Based on a comparison with the summed results from having run the analysis separately for each of the 20 species, the best performing composite network showed very similar prioritization of habitats (Spearman’s r = 0.976). This composite network was calculated based on the sum of the intra-patch connectivity for nodes and the average of the link probabilities weighted by the average suitable habitat area, requiring a quarter of the computing resources as the full species analysis representingan important gain in efficiency.

#### iv) Multi-node connectivity metrics

Connectivity analyses typically rank the importance of individual habitat patches in the network using single-node metrics from graph theory, e.g., probability of connectivity (PC) or integrated index of connectivity (IIC). Pereira et al. (2017) argue that, depending on their spatial arrangement, complementarity or redundancy, some groups of patches may better contribute to connectivity than the top individual patches. Through a study of 20 bird species in the Natura 2000 conservation network in Catalonia, Pereira and colleagues illustrate how two multi-node centrality metrics, ‘m-reach-closeness’ and ‘m-fragmentation’, drawn from social network theory (An and Liu 2016) are complementary and can be differentially employed depending on the movement capacity of each species. The m-reach-closeness metric identifies the set of nodes that is maximally connected to all other nodes, thereby prioritizing access across the entire network for high mobility species. In contrast, the m-fragmentation metric seeks to identify key patches that bridge core habitats, important for reducing species fragmentation in isolated populations with low mobility.6

#### v) Metapopulation capacity

Increasingly, connectivity analyses report the metapopulation capacity metric *lambda* of their final network prioritization. From metapopulation theory, the metapopulation capacity predicts the persistence of a population for a given landscape configuration based on rates of colonization and extinction (Ovaskainen and Hanski 2001). Based on an adjacency matrix, *lambda* is calculated as the leading eigenvalue of this matrix, where values above 1 indicate species persistence. In connectivity analyses it can be interpreted as the viability of a species population for a given habitat network configuration. In their study of 30 terrestrial mammals in Borneo, Brodie et al. (2016) apply *lambda* to identify network typologies that best support the community of species considered, i.e., the persistence of all species in the regional pool. Brodie et al. (2016) argue that an advantage of this approach is that it ranks links in the network according to their strength rather their presence-absence as with graph theory metrics. Furthermore, their final response variable (metacommunity stability) ranks these linkages based on the ultimate measure of interest, species persistence, rather than the proximate goal of network connectivity (Brodie et al. 2016).

#### vi) Multispecies connectivity optimization

Linear programming can identify an optimal network configuration that simultaneously meets habitat and connectivity requirements for two or more species. Due to their computation complexity, linear mixed integer programs traditionally include a single spatial attribute (connectivity) in their model resulting in long thin reserve designs that are likely suboptimal for species (Conrad et al. 2012). Wang and Önal (2016) design a linear integer programming model for multiple species that incorporates compactness in addition to the connectivity of landscape reserves. They apply their method to 10 bird species in Illinois to identify optimal reserves based on a minimum probability threshold. Their method also identifies multiple sub-reserves when a single reserve is inadequate for the overall species conservation goal. The authors explain that this is important when designing reserves for multiple species where habitats are scattered throughout the potential conservation area. In such cases, the spatial coherence of selected sites must be species-specific.

## 4. Discussion and Future Directions

The field of multispecies connectivity analysis has grown steadily over the past decade. This has led to a flourishing of approaches as scientists and conservation planners seek more ecologically effective and efficient methods to model multiple species habitat and movement needs. However, work is needed to better define the goals of multispecies connectivity analyses, to agree on metrics that evaluate and compare the performance of conservation designs, and most crucially to incorporate species interactions into network selection.

### 4.1 Setting a common definition and goals for MSC analyses

In this review we propose multispecies connectivity analysis is a ‘methodology for identifying a network of habitats and movement pathways that supports *the long-term persistence of multiple species in a landscape*’. Most papers we reviewed did not explicitly state species persistence as the goal of the analysis, but rather the identification of common habitat networks and movement corridors. While a laudable goal, corridors alone will not ensure species survival in landscapes if a minimum area of habitat is not also protected. Habitat area and connectivity must be assessed together. A small number of studies estimated the metapopulation capacity of prioritized networks *post hoc* to assess whether they support the persistence of species, with a smaller number using the metric to inform network design (e.g. Drielsma and Ferrier 2009; Brodie et al. 2015). The metapopulation capacity metric can be used to test the resilience of different network configurations under future land use and climate change scenarios (e.g. Shen et al. 2015). We believe this is an important innovation to advance the field by both providing a metric to rank potential network configurations across species and a robust means of comparing the effectiveness of alternative conservation network plans (Grantham et al. 2010).

Connectivity maps are sensitive to the way in which connectivity is formalized and implemented in models (Reed et al. 2017; Albert et al. 2017) and there is little consensus as to the effectiveness of different approaches (Marrec et al. 2020). As new techniques are devised to reduce data and processing requirements, more studies are needed to understand the trade-offs in time, data, accuracy and effectiveness that these approaches engender (e.g., Meurant et al. 2018; Jennings et al. 2020). With an increase in MSC approaches, developing common criteria and metrics will be vital to selecting from competing network designs and establishing best practices as the field of MSC modelling continues to grow Some studies are already tackling this challenge by using scenario-based simulation and common metrics to compare the effectiveness of different MSC approaches under climate change (Rayfield et al. *in prep*).

### 4.2 Need for greater network validation

Across studies there was a striking lack of empirical validation of multiple species connectivity models. Only a handful of reviewed papers used independent datasets to validate the accuracy of their networks or select amongst them (see Koen et al. 2014; Marrotte et al. 2017; Brennan et al. 2020). Of these, most relied on genetic data to assess how habitat fragmentation and corridors influence functional connectivity. Without validation, it is not possible to determine how effective different MSC methods are for predicting and conserving species connectivity. For instance, Marrotte et al. (2017) compared node-based estimates of genetic connectivity using neutral microsatellites for a set of mammals across Ontario with modelled estimates of current density. They found that current density was proportional to the probability of movement in fragmented parts of the landscape, but not where habitat was abundant. Furthermore, in their model high current density did not reflect high gene flow, rather, it identified pinch points restricting species movements. Using a naturalness-based approach, Koen et al. (2014) found that modelled current density was strongly correlated with between empirical roadkill and fisher movement patterns. As movement and occupancy data become more readily available through less expensive genetic sequencing and open data repositories, the validation of connectivity results should become a standard part of robust MSC analyses.

### 4.3 Integrating species interactions

Despite the novel methods noted in this review, we find that the field is largely in a nascent stage with respect to its ability to meaningfully incorporate multiple species interactions into connectivity modelling. None of the studies reviewed incorporated behavioral or population dynamics between co-occurring species (but see Shahnaseri et al. 2019 who included prey abundance in their habitat distribution model for focal predators). Studies continue to “stack” independent species networks to prioritize corridors rather than building “multilayer networks” that include ecological dependencies and interactions across layers (Kéfi et al. 2016; Pilosof et al. 2017). This omission in MSC analyses is critical as functional connectivity is not only shaped by landscape structure, but species interactions as well (Gonzalez, Rayfield, and Lindo 2011; Courbin et al. 2014). For example, in a study of a wolf-moose-caribou predator-prey system in a fragmented landscape of Quebec, Courbin et al. (2014) found that wolves use indicators of prey habitat quality and preference, rather than the distribution of prey *per se*, to orient their movement in landscapes where prey are highly mobile. Such studies illustrate that to accurately predict movement in landscapes, MSC analyses must integrate metacommunity approaches that consider food-webs and the spatial dynamics of interacting species into modelling approaches (e.g., Yeakel et al. 2020).

One MSC analysis that did incorporate spatial species dynamics is Rayfield et al. (2009), a study published prior our review horizon. In it, Rayfield and colleagues develop a general framework to incorporate consumer-resources dynamics into spatial conservation networks. Their approach protects areas that maintain the connectivity between the distribution of consumers by using an interaction kernel that defines the probability distribution of foraging distances based on the movement abilities of the consumers and resources. When applied to the case of the American marten (*Martes americana)* and its prey (the red-backed vole, *Myodes rutilus*, and the deer mouse, *Peromyscus maniculatus*), their method prioritized spatially aggregated reserves that maintain local habitat quality for all species. Similarly, using a theoretical framework, Baggio et al. (2011) developed an agent-based model to explore connectivity designs while considering spatial predator-prey interactions in fragmented landscapes. Their model also concluded that both predator and prey benefit most from globally well-connected habitat patches. Results from both studies aligned with empirical findings of Courbin et al. (2014) on the wolf-moose-caribou system in which prey selected habitat patches that were connected by multiple links instead of isolated ones. The authors suggest that predators are cued into these connectivity preferences.

Advancements emerging from food-web modelling to prioritize habitat conservation may provide new tools to better incorporate multispecies interactions into connectivity assessments (Yeakel et al. 2020). Promising studies on metawebs suggest that interacting species can be meaningfully grouped into trophic guilds based on species interactions and functional traits to understand spatial variation of food webs (O’Connor et al. 2020). Such studies can help bridge the gap between spatial community ecology and landscape ecology, a link that is currently missing in applied connectivity conservation.

## 5. Conclusions

This review showed the breadth of methods available to analyze and prioritize multispecies connectivity. It also revealed that more work is needed to test and validate different approaches across a common set of criteria to establish a set of best practices to inform conservation planning. To do this, more comparative studies that contrast methods within landscapes are needed to test efficiency and accuracy. This research would be further strengthened by increased analyses of uncertainty and sensitivity to better understand which steps in MSC modelling should we invest in to reduce uncertainty. Finally, the development and expansion of observation and monitoring networks will be key to provide timeseries data to validate and update MSC analyses through high-resolution real-time data on species movement patterns in dynamic and evolving landscapes.

## 6. Authors’ contributions

AG, BR, CP, AEM, SW contributed equally to the conception and design of the study and to final review of the manuscript. SLRW, OT, FM, VDL, KMTL, ASD all contributed equally to the review the literature, data collection and revision of the manuscript. SLRW was responsible for data analysis and writing of the manuscript.

## 7. Data Availability

Data from the literature review will be made available on the Open Science Foundation’s repository (upon publication). The full list of papers included in the review and references will be provided therein.

## 8. Acknowledgements

We acknowledge the following conflict of interest. The work is the product of a contract from Environment and Climate Change Canada to Habitat and ApexRMS (private companies) to characterize the current state of multi-species connectivity science. The following authors are employed by Habitat SLRW, KMTL, VDL, ASD, OT, FM, and AG is a co-founder of Habitat. The following authors are employed by ApexRMS, BR.

